# Moving patronin foci and growing microtubule plus ends direct the spatiotemporal dynamics of Rho signaling and myosin during apical constriction

**DOI:** 10.1101/2021.01.08.425813

**Authors:** Anwesha Guru, Surat Saravanan, Deepanshu Sharma, Maithreyi Narasimha

**Author notes:** equal contributions (Anwesha Guru, Surat Saravanan).

## Abstract

The contraction of the amnioserosa by apical constriction provides the major force for Drosophila dorsal closure. The nucleation, movement and dispersal of apicomedial actomyosin complexes generate pulsed constrictions during early dorsal closure whereas persistent apicomedial and circumapical actomyosin complexes drive the unpulsed constrictions that follow. What governs the spatiotemporal assembly of these distinct complexes, endows them with their pulsatile dynamics, and directs their motility remains unresolved. Here we identify an essential role for microtubule growth in regulating the timely contraction of the amnioserosa. We show that a symmetric cage of apical microtubules forms around the coalescing apicomedial myosin complex. An asymmetric tail of microtubules then trails the moving myosin complex and disperses as the myosin complex dissolves. Perturbing microtubule growth reduced the coalescence and movement of apicomedial myosin complexes and redistributed myosin and its activator, Rho kinase to the circumapical pool and altered the cell constriction and tissue contraction dynamics of the amnioserosa. We show that RhoGEF2, the activator of the Rho1 GTPase, is transiently associated with microtubule plus end binding protein EB1 and the apicomedial actomyosin complex. Our results suggest that microtubule growth from moving patronin platforms modulates actomyosin contractility through the spatiotemporal regulation of Rho1 activity. We propose that microtubule reorganisation enables a self-organising, mechanosensitive feedback loop that buffers the tissue against mechanical stresses by modulating actomyosin contractility.

## Introduction

Apical constriction powers a wide array of morphogenetic movements including tissue invagination, tissue contraction and single cell behaviours such as cell delamination or extrusion. Much work on Drosophila ventral furrow invagination has informed our understanding about the subcellular mechanisms that accomplish apical constriction and the signalling pathways that regulate them. These studies have demonstrated the requirement for the regulation of actomyosin cytoskeleton dynamics by transcription factor dependent activation of GPCR signalling, culminating in the activation of the small GTPase Rho (Kerridge et al., 2016; Kolsch et al., 2007; Martin et al., 2009; Sweeton et al., 1991). They have also highlighted the importance of biomechanical feedback and the differential, radially polarised regulation of Rho GTPase activation in the medial and circumapical actomyosin pools for achieving pulsed constriction (Mason et al., 2013; Mason et al., 2016; Munjal et al., 2015). What governs the differential subcellular distributions of the Rho activators and effectors in this context remains unclear.

Apical constriction also powers the contraction of the amnioserosa, the major force provider for dorsal closure. Unlike apical constriction during ventral furrow formation in which constrictions are racheted with each pulse made up of a constriction phase and a stabilisation phase during which shape change is irreversible, apical constriction in the amnioserosa is characterised by an early phase (Phase I) during which unracheted constriction results in high amplitude pulses of constriction and relaxation that generate little or no net reduction in apical area. This is followed by a phase (Phase II) of dampened pulses and net reduction in area. Phase I is mediated by motile, medial, actomyosin complexes that cause local, anisotropic constriction. Phase II relies on junctional and persistent apicomedial actin networks. In addition, a small fraction of amnioserosa cells constrict their apices rapidly and extrude or delaminate. This behaviour requires differential actomyosin contractility in the delaminating cell and its nearest neighbours, and is associated with pulse dampening in both populations (Blanchard et al., 2010; Dehapiot et al., 2020; Saravanan et al., 2013). Although the consequences of RhoGEF2 downregulation in the amnioserosa underscore the involvement of Rho GTPase signalling, the upstream regulators that activate distinct constriction programs in the amnioserosa remain unclear (Azevedo et al., 2011). Earlier work from our lab revealed that anisotropies in interfacial tension generated either by subcellular laser ablation or by activating Rho in a single amnioserosa cell was sufficient to direct the movement of medial actomyosin complexes in the neighbours towards the interfaces with the ablated cell or the cell in which Rho was activated (Meghana et al., 2011; Saravanan et al., 2013). How these anisotropies are generated and sensed to result in actomyosin complex movement remains poorly understood.

The microtubule cytoskeleton enables the transport of proteins and organelles to distinct subcellular locations through the action of motor proteins. In addition, microtubules confer mechanical integrity to cells by enabling them to resist compression, and can generate force against a membrane either by polymerising or depolymerising behind it, or by being anchored to the cell cortex (Brouhard and Rice, 2018). Microtubules govern cell shape in plant and animal cells and can be reorganised in response to it (Bidhendi et al., 2019; Chakrabortty et al., 2018; Gomez et al., 2016; Mirabet et al., 2018). The integrity of the microtubule cytoskeleton has been shown to be necessary for apical constriction during Xenopus gastrulation, vertebrate neuronal delamination, and during Drosophila salivary gland and eye imaginal disc morphogenesis (Booth et al., 2014; Fernandes et al., 2014; Kasioulis et al., 2017; Lee et al., 2007).

Previous work from our lab uncovered the striking reorganisation of the apical microtubule meshwork in amnioserosa cells during native and laser-induced delamination. This reorganisation was characterised by the alignment and polarisation of microtubules in the apical plane in the nearest neighbours of a delaminating cell which also exhibited shape transformations, and provided evidence for their dynamic nature. While disrupting microtubule integrity by microtubule severing altered the dynamics of delamination and delayed cell extrusion, whether and how apical microtubules contribute to apical constriction in the amnioserosa remains unclear (Meghana et al., 2011). In the work presented here, we use a combination of genetic perturbations that affect microtubule organisation and dynamics targeted to amnioserosa cells and high-resolution live confocal microscopy to examine the role of the microtubule cytoskeleton for amnioserosa contraction during Drosophila dorsal closure. Our results uncover the importance of persistent microtubule growth in governing the organisation and dynamics of actomyosin necessary for timely tissue contraction, and identify mechanistic links between the actomyosin and microtubule cytoskeleton.

## Results

During Phase I of Drosophila dorsal closure, high amplitude pulses of constriction and relaxation generate anisotropic cell shape changes in the amnioserosa with little or no net reduction in apical area. Phase II which follows is characterised by dampened pulses and net isotropic reduction in area. Earlier work from many labs including ours has uncovered the distinct localisation and dynamics of apicomedial myosin associated with each phase (Blanchard et al., 2010; Saravanan et al., 2013; Solon et al., 2009). During Phase I, each myosin cycle (visualised using Sqh GFP, in which GFP is fused to the regulatory light chain of non-muscle myosin) begins with the nucleation (Figure S2 A1, A2) of an apicomedial myosin ‘seed’ that grows by coalescence and aggregation to form a larger ‘blob’ (Figure S2 A2, A3). This blob moves in the apical plane for a distance, towards an interface and constricts the apical membrane locally (Figure S2 A4-A6) before it dissolves and disappears (Figure S2 A7). After a lag phase of approximately 1 minute, a new cycle commences. Coalescence and movement can happen simultaneously in some cycles. An average myosin cycle takes approximately 2 mins [115+/− 17 secs; n=40; (Blanchard et al., 2010; Saravanan et al., 2013; Solon et al., 2009)]. The formation-dissolution cycle of myosin temporally correlates with the constriction and expansion phases of a single pulse, with the lowest apical cell surface area associated with the presence of the most intense and tightly coalesced myosin blob (Saravanan et al., 2013). Phase II is associated with circumapical myosin enrichment and non-motile and persistent apicomedial actomyosin networks (Blanchard et al., 2010; Dehapiot et al., 2020; Saravanan et al., 2013). What directs the spatiotemporal attributes of actomyosin dynamics, and in particular its movement remained unclear. This motivated us to examine the influence of the microtubule cytoskeleton on amnioserosa cell constriction and tissue contraction.

### Moving microtubule platforms associate with apicomedial myosin during pulsed apical constriction

The microtubule cytoskeleton whose dynamics is governed by microtubule associated proteins interacting with its ends, provides platforms that can integrate signalling and mechanics (Brouhard and Rice, 2018). Epithelial cells are characterised by at least three microtubule pools: an apical meshwork, lateral arrays running apicobasally along the cell cortex or located medially, and a basal mat (Bacallao et al., 1989; Bartolini and Gundersen, 2006; Toya and Takeichi, 2016). This was also found to be the case in the amnioserosa cells. The apical meshwork forms an array of seemingly unaligned, randomly oriented microtubules that do not show any characteristic association with the centrosomes (Figure S1 A, B). The lateral pool forms a parallel array of aligned microtubules along the apicobasal axis. The basal pool forms a randomly oriented array /carpet (Figure S1 A). Real time analysis of microtubule organisation using Jupiter-GFP [a GFP fusion protein that labels microtubules along its entire length (Karpova et al., 2006)] uncovered the dynamics of the apical meshwork. Single filaments exhibited buckling and length changes. In addition, microtubule collectives exhibited two kinds of configurations that correlated with the pulse cycle. Long microtubules or microtubule bundles exhibited “bundling” and splaying that were associated respectively, with the constriction and relaxation phases of a pulse. Microtubules aligned along the major axis of constriction and were brought into closer proximity during “bundling” and dispersed and adopted random orientations during splaying (Figure S2 F, G and Video S2). A subset of short microtubules organised into aster/cage-like configurations (Figure S2E, Video S2). Live imaging with Jupiter GFP and ECadherin GFP revealed that both aster/cage formation and bundling were temporally correlated with the constriction phase of each pulse (Figure S2 E, G).

To determine whether the dynamic microtubule reorganisations we observed were polarised, and to establish the nature of its association with apicomedial myosin pools, we examined the dynamics of microtubule ends by visualising the dynamics of the plus and minus end associated proteins EB1 and Patronin that respectively facilitate and suppress dynamics at the microtubule ends (Goodwin and Vale, 2010; Akhmanova and Steinmetz, 2015), using Ubiquitin promoter driven Patronin GFP and genomic EB1 GFP fusion protein expressing transgenes (Bulgakova et al, 2013; Wang et al., 2013) alongside myosin (visualised using Sqh mCherry). A large patronin cloud prefigured the appearance of the myosin seed, and condensed symmetrically around it as the seed grew and coalesced. During the movement of the myosin complex, patronin foci formed a moving platform that trailed the myosin blob some distance away from it. As the blob fragmented and the cell relaxed, the patronin foci moved centrifugally to form a dispersed cloud again (Figures 1A, S2B and Video S1A). The distinct organisation of the patronin cloud combined with the reduced dynamics of the minus ends made it possible to quantitatively assess the relationship between myosin and microtubules. A myosin seed always appeared within the patronin cloud, which condensed around it (100%, n=51 blobs from 19 embryos).

**Figure 1:**
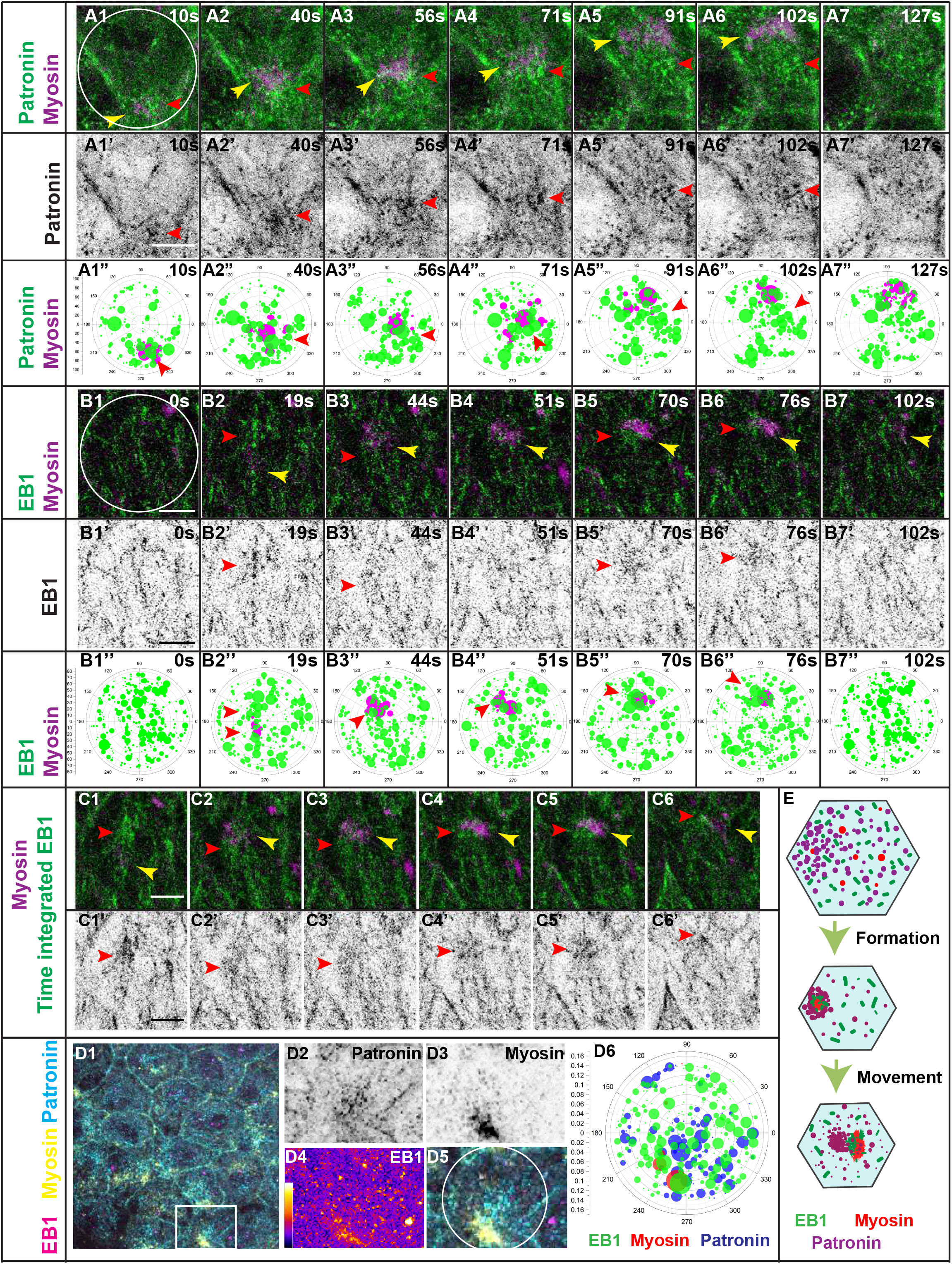
Moving microtubule platforms associate with apicomedial myosin during pulsed apical constriction. (A1-A7) Time-lapse images of an amnioserosa cell in Phase I of dorsal closure from embryos carrying Patronin GFP (green) and Sqh mCherry (magenta), showing compaction of the patronin cloud (green) around the coalescing myosin blob and then trailing it during its movement. A1’-A7’ are the corresponding single channel images of Patronin. A1”-A7” are the corresponding polar plots made using the ROI marked in A1 for Patronin (green) and Myosin (magenta). (B1-B7) Time-lapse images of an amnioserosa cell expressing EB1 GFP (green) and Sqh mCherry (magenta) during pulsed apical constriction. B1’-B7’ are the corresponding single channel images of EB1 GFP. B1”-B7” are the corresponding polar plots made using the ROI marked in B1 for EB1 (green) and Myosin (magenta). (C1-C6) Time lapse images of the pulse cycle shown in B, with Sqh mCherry (magenta) and time integrated EB1 GFP tracks (green). Each frame is an integration of three successive frames. (C1’-C6’) are the corresponding single channel images of time integrated EB1 GFP. Yellow arrowhead indicates the myosin blob. Red arrowheads point towards the reorganisation of EB1 (B, C) or Patronin (A) Scale bar-5μm. (D1-D6) Merge (D1) of an apical projection of the amnioserosa of a Patronin GFP embryo stained for GFP (cyan), pMLC (yellow), and EB1 (magenta). D2-D5 are the cropped single channel and merge images of a single myosin blob highlighted with a white box in D1. (D6) Polar plot made using the ROI marked in D5 showing spatial distribution of myosin (red), Patronin (blue) and EB1 (green). (E) Schematics describing the organisation of microtubule ends in relation to the formation and movement of a myosin blob during pulsed apical constriction. (See also Figure S2 and Video S1.)

A subset of EB1 comets reorganised to form a transient, close association with apicomedial myosin in the form of an aster/cage as the cell constricted (Figure S2 C, D). This association was often evident as soon as a seed of myosin was visible and the EB1 comets were initially symmetrically distributed around it as it grew to form a blob. As the blob moved in the apical plane, the comets were asymmetrically distributed and trailed the myosin blob while maintaining close proximity to it. New comets became associated with it during coalescence and/movement and the comets retracted away from the apicomedial myosin complex as it dissolved (Figure 1 B, C and Video S1 B, C).

Polar plots also captured the dynamic associations of the microtubule ends with myosin described above (Figure 1 A”, B”). Triple labelling of amnioserosa cells with EB1, patronin and myosin further strengthened their relative radial positions. Myosin formed the central core, with microtubule plus ends tightly congregated in close proximity to it and the minus ends farther away and distributed over a larger area (Figure 1D). These results suggest that a subset of short, dynamic microtubules surround the myosin blob to form a predominantly “minus-end out” cage during its coalescence and then redistribute asymmetrically around it, to form a tail emanating from a patronin platform, with EB1 comets in close proximity with it (Figure 1E). Technical limitations and the presence of a small fraction of patronin in close proximity to myosin do not allow us to exclude the possibility that a subset of microtubules organise into antiparallel bundles. These observations raise the possibility that the microtubule cytoskeleton might regulate the organisation or dynamics of myosin during pulsed apical constriction and through it, influence cell and tissue dynamics.

### Perturbing microtubule growth alters amnioserosa cell constriction and contraction dynamics

To test whether microtubule growth influences cell and tissue dynamics in the amnioserosa during dorsal closure, we drove the expression of a truncated version of EB1 previously shown to act as a dominant negative (EB1-DN) and to reduce microtubule growth duration without affect growth velocity in both mammalian cells and Drosophila tissues (Komarova et al., 2009; Bulgakova et al., 2013), using an amnioserosa specific Gal4 driver. In addition, we found that expression of UAS EB1-DN did not grossly alter the architecture of the apical microtubule meshwork (Figure S1 D, G).

We used ECadherin GFP in the background to allow quantitative morphodynamic analysis of the amnioserosa. Significant reductions in pulse amplitude were observed in Phase I upon expression of EB1-DN (Figure 2A, see Methods). A similar reduction in pulse amplitude was also observed when the microtubule meshwork was disrupted by the overexpression of the severing protein Spastin, demonstrating the necessity for an intact meshwork, but not upon the overexpression of Tau, a potential regulator of microtubule stability (Figure S3I). Both these perturbations also starkly altered microtubule architecture, with the former reducing microtubule length and mass and the latter causing pronounced bundling and length changes (Figure S1 D-F). The basis for their effects is therefore likely to be more global or complex. To determine whether microtubule growth affects contractility, we examined the real time apical area dynamics of control and EB1-DN expressing amnioserosa cells (see Methods). Such an analysis revealed that although the mean area of EB1-DN expressing cells is marginally higher than that of wildtype cells during Phase I, their constriction rate was higher during early closure but comparable towards the end of closure (Figure 2 B, C). This, combined with the reduction in pulse amplitude observed in EB1-DN cells (Figure 2 A), suggests that perturbing persistent microtubule growth may lead to the precocious initiation of rapid and unpulsed constriction.

**Figure 2:**
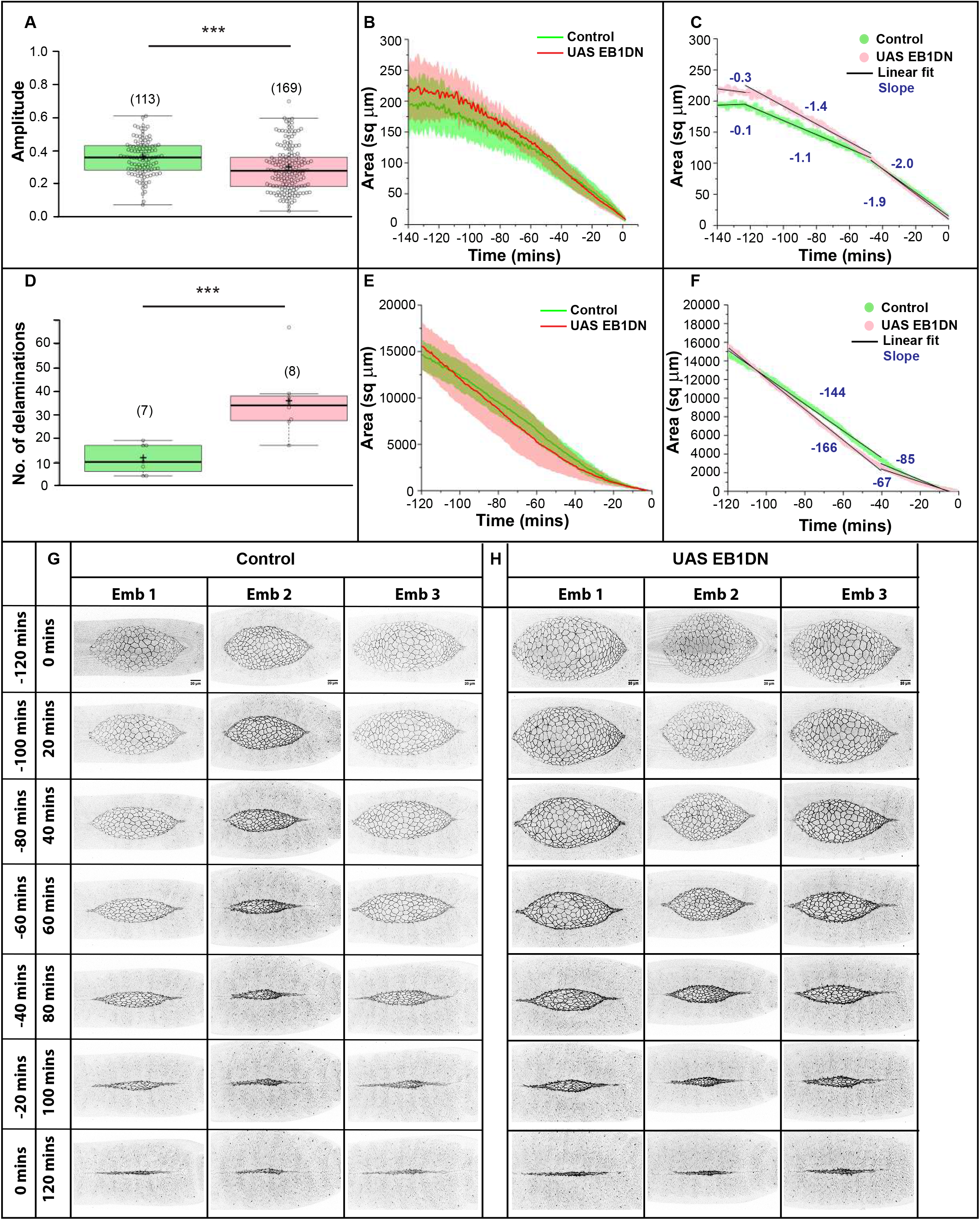
Persistent microtubule growth influences cell and tissue dynamics in the amnioserosa. (A) Normalised pulse amplitude of amnioserosa cells from control embryos (green) and embryos overexpressing EB1-DN (pink) in the amnioserosa during phase I of dorsal closure (see Methods). (B) Apical cell area dynamics (mean ± sd) of amnioserosa cells from embryos overexpressing EB1-DN (n=34 cells from 7 embryos, pink) and controls (n=44 cells from 9 embryos, green) over a period of 140 minutes. (C) Mean apical area dynamics of amnioserosa cells in EB1-DN (pink dots) and control (green dots) with the corresponding linear regression line fits for each of the three phases of dorsal closure. The slopes of fitted lines for each of the phases in both genotypes are indicated. (D) Number of delamination events in control embryos (green) and embryos overexpressing EB1-DN (pink) (see Methods). (E) Area dynamics (mean ± sd) of the amnioserosa from embryos overexpressing EB1-DN (n=6 embryos, pink) and controls (n=6 embryos, green) over a period of 120 minutes. (F) Mean area dynamics of amnioserosa in EB1-DN (pink dots) and control (green dots) with the corresponding linear regression line fits and the corresponding slopes are indicated. (G-H) Time-lapse images of maximum intensity projections of embryos expressing ECadherin GFP without (G) or with the overexpression of EB1-DN (H) in the amnioserosa. In the box plots in A and C, boxes show median (dark line) ± interquartile range. The mean is also indicated by ‘+’. The sample size is given in brackets. ***-p<0.0001. (See also Video S4B.)

The temporal dynamics of amnioserosa contraction was also affected. A retrospective analysis of area dynamics revealed that the area of the amnioserosa during early closure was often larger and more variable in EB1-DN embryos compared to controls but were indistinguishable from wildtype in the latter half. Surprisingly the rate of contraction during early closure was significantly higher in EB1-DN embryos compared to controls (Figure 2 E-H & Video S4B). These differences mirror the differences in cell area dynamics and suggest that differences in cell constriction dynamics might contribute to the differences in contraction dynamics. In addition, the frequency of cell delamination - a rare and stochastic cell behaviour associated with rapid apical constriction and cell extrusion - (Meghana et al., 2011, Muliyil et al., 2011), was higher in EB1-DN embryos compared to controls (Figure 2D). These results demonstrate the influence of microtubule growth in orchestrating cell and tissue dynamics during dorsal closure. Whether an increase in the constriction rate predisposes cells to delamination either cell autonomously or through its effects on tissue tension, or whether the increase in frequency of delamination contributes to the early increase in contraction rate (Toyama et al., 2008; Muliyil and Narasimha, 2014) remains unresolved. To address the mechanisms that might underlie the differences in pulse amplitude and constriction rates upon perturbing microtubule growth, and given the close association between apicomedial myosin and microtubules, we examined its effects on the distribution and dynamics of myosin.

### Microtubule growth modulates the organisation, subcellular distribution and movement of apicomedial myosin

In control embryos, apicomedial myosin exhibited the pulsed dynamics described previously (Figures S1 A and 3A and Video S4A). In cells overexpressing the EB1-DN, apicomedial myosin foci failed to coalesce into a compact blob and appeared either as multiple distinct punctae or were not visible at all (Figure 3B). In addition, the circumapical pool of myosin was more prominent compared to controls. Perturbing microtubule growth did not significantly influence the formation, dissolution and total persistence times of medial myosin (see Methods). However, myosin path length, which we define as the distance travelled by the medial myosin blob prior to its dissolution was significantly lower (Figure 3 C-G and Video S4A). Spastin and Tau overexpression also influenced myosin coalescence and path length, albeit in opposite directions, with the former reducing and the latter increasing it, presumably because of their effects on microtubule length and possibly also stability (Figure S3, A-C, G-H). The effects of EB1-DN however clearly demonstrate the importance of regulating microtubule growth persistence on myosin coalescence and movement and in regulating the distribution of myosin between circumapical and apicomedial pools.

**Figure 3:**
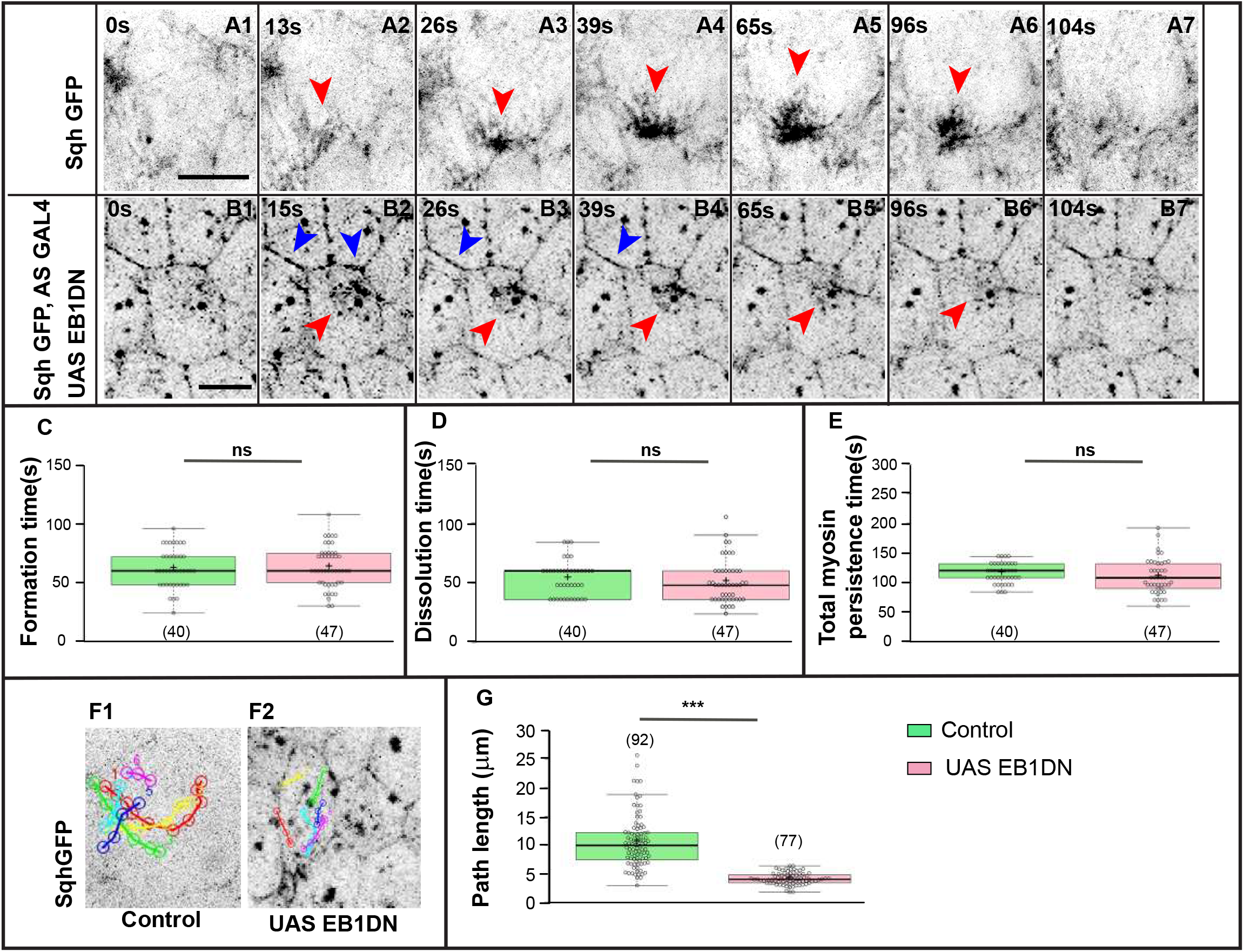
Persistent microtubule growth influences apicomedial myosin organisation and dynamics. (A-B) Time-lapse images of single amnioserosa cells during Phase I of dorsal closure showing qualitative changes in the spatial organisation of apicomedial myosin (visualised using Sqh GFP) in embryos that are otherwise wild type ([A1-A7; (n=40 myosin cycles from 4 embryos)], or overexpress EB1-DN [B1-B7; (n=47 myosin cycles from 5 embryos)] in the amnioserosa. Red arrowheads point to apicomedial myosin complexes. Blue arrowheads indicate the enriched circumapical myosin pool. Scale bar −10μm. (C-E) Myosin cycle times (formation-C, dissolution-D and total myosin persistence times-E) measured in wildtype embryos, and in embryos overexpressing EB1-DN in a Sqh GFP /+ background. (F) Representative tracks of medial myosin movement in a single Phase I amnioserosa cell from control embryos (F1), and from embryos overexpressing EB1-DN (F2). Each track represents the distance travelled by one apicomedial myosin blob/structure in one cycle. Many myosin cycles (colour coded) were tracked in each cell. (G) Quantitative analysis of apicomedial myosin path lengths in each of the genotypes analysed. In the Box plots in C-E, G, boxes show median (dark line) ± interquartile range. The mean is also indicated by ‘+’. The sample size is given in brackets. ***-p<0.0001. (See also Figure S3 and Video S4A.)

### EB1-RhoGEF2 interactions govern the spatial distribution of Rho pathway activity

We reasoned that microtubule growth might modulate myosin distribution, coalescence and movement either by aiding myosin transport and/or by controlling the delivery of regulators of actomyosin contractility. The velocity of motile apicomedial myosin complexes, inferred from tracks of the kind shown in Figure 3 F and Video S4A, was found to be 0.16 μm/sec (± 0.04; n=121 tracks from 4 embryos). The velocity of microtubule growth inferred from the displacement of EB1 comets in movies of the kind shown in Video S3 was 0.25 ± 0.08 μm/sec (n=101 punctae, N=4 embryos). This comparison shows that the observed velocity of apicomedial myosin is close to the velocity of microtubule growth (0.1 μm/sec) and suggests that microtubules might modulate myosin by mechanisms independent of their ability to serve as tracks for myosin transport. One possibility that is consistent with the observed speeds is that proteins associated with the growing ends of microtubules may regulate actomyosin contractility and myosin movement. One candidate protein that fulfils both criteria is RhoGEF2, which activates Rho and has also been shown to bind microtubule plus ends in an EB1 dependent manner (Rogers et al., 2004). Live imaging of RhoGEF2 GFP revealed its pulsatile dynamics, characterised by the centripetal movement and condensation of RhoGEF2 nodes upon cell constriction, and its centrifugal dispersal upon cell relaxation (Figure 4A, Video S5A). The dynamics of RhoGEF2 nodes was reminiscent of the dynamics of EB1 comets (Video S3). The measured mean velocity of RhoGEF2 punctae (0.27 ± 0.07 μm/sec, n=96 punctae from 5 embryos) from live images was also remarkably similar to the mean velocity of EB1 comets (0.25 ± 0.08 μm/sec n=101 punctae from 4 embryos). In fixed preparations immunostained for RhoGEF2 GFP, apicomedial RhoGEF2 nodes were found in some cells (presumably in the constriction phase) to be closely associated with the actomyosin blob, with a subset of microtubules that appeared to be aligned towards it (Figure 4B).

**Figure 4:**
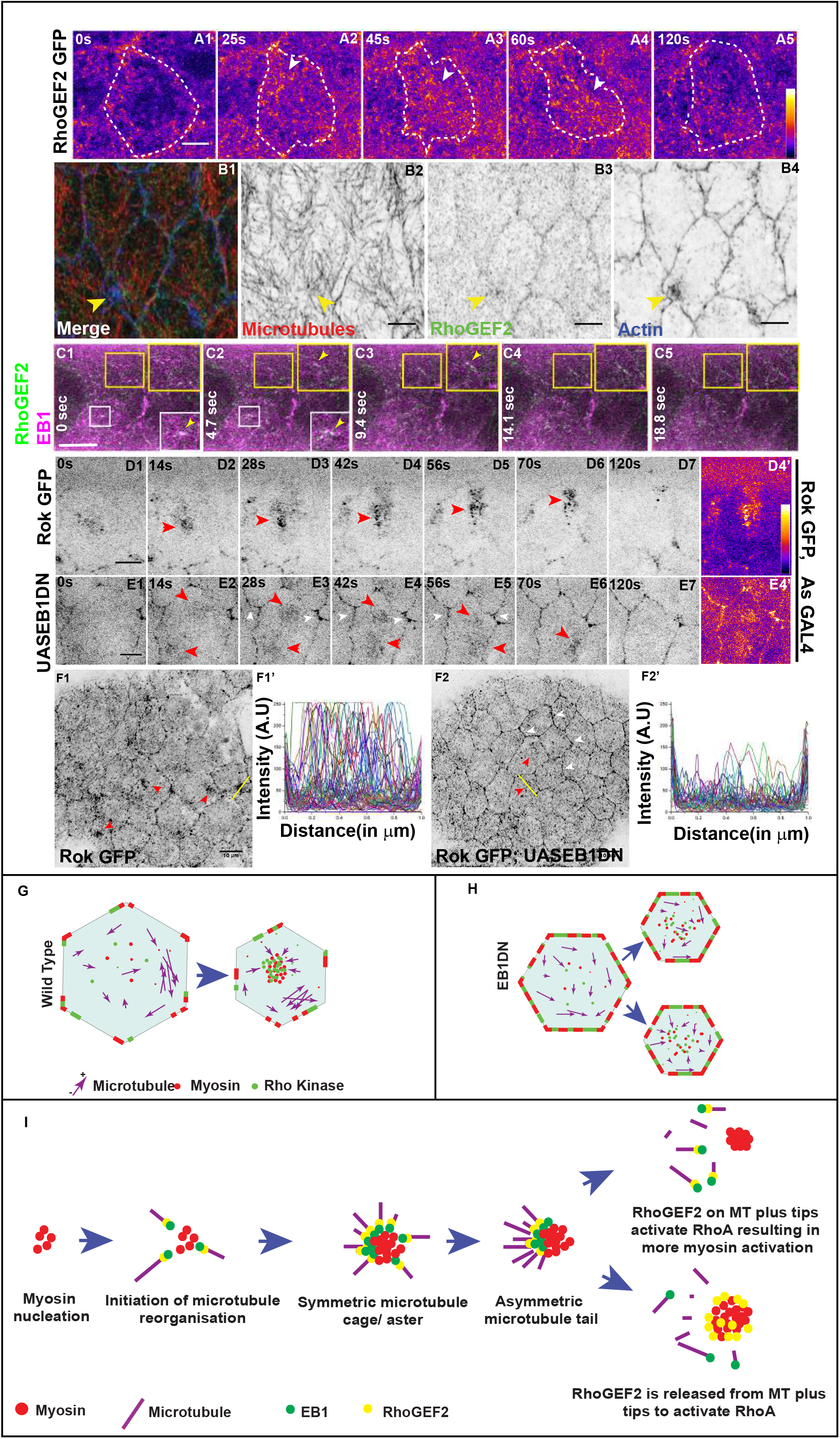
EB1-RhoGEF2 interactions govern the spatial distribution of Rho pathway activity. (A1-A5) Intensity coded time-lapse images of amnioserosa cells in Phase I of dorsal closure, from an embryo carrying RhoGEF2 GFP. The dashed white lines mark cell outlines and the white arrowheads point to the congregation and dispersal dynamics of apicomedial RhoGEF2 GFP clusters. Scale bar-5 μm. (B1-B4) Merge (B1) and single channel images (B2-B4) of an apical projection of the amnioserosa of a RhoGEF2 GFP embryo stained for alpha tubulin (red, grey in B2), GFP (green, grey in B3) and phalloidin (blue, grey in B4). Yellow arrowheads point to the spatial proximity of medial actin and RhoGEF2 clusters and microtubule reorganisation. Scale bar-5 μm. (C1-C5) Time lapse images showing colocalization (white) of moving EB1 comets (magenta) and RhoGEF2 (green). Yellow and white insets are magnified views of the areas contained within the respective boxes. Yellow arrowheads points to the colocalization. (D, E) Time-lapse images of maximum intensity projections of apical slices of amnioserosa cells from Phase I of dorsal closure, from embryos carrying Rok GFP in otherwise wildtype embryos (D1-D7) and in embryos overexpressing EB1-DN in the amnioserosa (E1-E7). Red arrowheads mark the apicomedial clusters of Rok GFP. White arrowheads indicate the enriched circumapical Rok pool. D4’ and E4’ are intensity coded fire images of D4 and E4 respectively. Scale bar-5 μm. (F1-F2) Maximum intensity projections of apical slices of amnioserosa cells from Phase I of dorsal closure from control (F1) and EB1-DN overexpressing embryos (F2) in which Rok GFP is visualized using immunofluorescence. Red arrowheads mark the apicomedial Rok clusters. (F1’, F2’) Line intensity profiles for Rok GFP in control (F1’, n=57 cells from 27 embryos) and EB1DN expressing cells (F2’, n=38 cells from 17 embryos). Representative yellow lines in F1, F2 show how intensity profiles in F1’ and F2’ were obtained (see Methods). Scale bar - 10 μm. (G, H) A schematic representation of the spatial distribution of myosin and its effector, Rok in the apicomedial region of a relaxed (left) or constricted (right) amnioserosa cell during Phase I of dorsal closure in a wild type (G) and upon EB1 DN overexpression (H). (I) A putative model for the temporal regulation of the interplay between the apicomedial myosin and the apical microtubule meshwork during pulsed constriction. (See also Video S5.)

To directly test whether RhoGEF2 is associated with microtubule plus ends, we simultaneously imaged the dynamics of EB1 (using UAS EB1 RFP expressed in the amnioserosa) and RhoGEF2 (using GFP transgene mentioned above) in real time. This allowed us to capture the transient association (for between 4 to 14 secs) of EB1 with Rho GEF2 and their subsequent dissociation (Figure 4C, Video S5B). These findings demonstrate that RhoGEF2 movement and subcellular distribution may depend on microtubule growth and shrinkage through its association with EB1, and provide an attractive mechanism by which the microtubule cytoskeleton can influence the spatial regulation of actomyosin contractility.

To determine whether the effect of perturbing persistent microtubule growth on myosin dynamics may rely on EB1-RhoGEF2 interaction dependent Rho activation, we examined the dynamics of Rho kinase/Rok (using a sqh promoter driven Rok GFP fusion protein), the key effector of actomyosin contractility downstream of Rho, in EB1-DN expressing central amnioserosa cells. Time-lapse imaging revealed that Rok exhibited pulsatile dynamics in wildtype cells. This was characterised by the centripetal movement and congregation of discrete Rok punctae to form a cluster when the cell constricted, and its dispersal by centrifugal movement as the cell relaxed, and was reminiscent of the behaviour of both EB1 comets and RhoGEF2 nodes (Videos S3 and S5). The brightest pixels of Rok GFP in the cell resided in the apicomedial cluster, whereas the circumapical regions contained sparsely distributed punctae of Rok (Figure 4D and Video S5C). This was also evident in fixed preparations in which Rok GFP intensities were quantified: substantially higher intensities of GFP were found in the apicomedial pool compared to the circumapical pool (Figure 4F1). When persistent microtubule growth was perturbed by EB1-DN, the apicomedial clusters of Rok appeared more dispersed and less bright, while Rok in the circumapical pool was brighter and more continuous (Figure 4E, 4F2 and Video S5C). These results demonstrate that persistent microtubule growth is required to maintain the spatial/subcellular distribution of Rok, specifically the formation of apicomedial clusters, and are consistent with its effects on the distribution of myosin. They uncover the requirement of RhoGEF2-EB1 interactions in regulating the spatiotemporal activation of Rho GTPase signalling during pulsed apical constriction and also provide an explanation for its effects on amnioserosa contraction (2379).

## Discussion

The work we describe uncovers for the first time, an essential requirement for microtubule growth in the regulation of apical constriction and tissue contraction during dorsal closure. It suggests that microtubule nucleation and growth from dynamically reorganising patronin foci provide moving platforms (Figure 1E), that modulate the spatiotemporal activation of actomyosin contractility through the local regulation of Rho GTPase activity. We propose that this in turn governs the force distribution between apicomedial and junctional pools of actomyosin and through it, the mode and rate of apical constriction (Figure 4 G-H). Our work suggests a model in which the rapid reorganisation of the apical microtubule meshwork in the apical plane can enable the mechanosensitive tuning of actomyosin contractility and uncovers a previously undescribed mechanism by which microtubules can influence apical constriction, tissue contraction and epithelial fusion morphogenesis (Booth et al., 2014; Ko et al., 2019; Jankovics and Brunner, 2006).

Two modes of apical microtubule meshwork reorganisation during pulsed constriction were identified in this study: long microtubules which exhibited alignment, bundling and splaying, and short microtubules that formed asters/cages and tails. We propose that the former provides a resistive network, possibly analogous to the recently described persistent actin network (Dehapiot et al, 2020), while the latter facilitates the local fine tuning of cell shape, dynamics and mechanics. In vitro reconstitution experiments and simulations have revealed that asters composed of short microtubules and nematic bundles composed of long extensible microtubules are observed in distinct phase spaces that are defined by ratios of microtubules to motor number or speed (Roostalu et al., 2018). The mechanisms that underlie the generation of the two microtubule configurations we observe in vivo, and the consequences of perturbing them individually will be interesting to investigate.

The experiments with spastin or tau overexpression establish the importance of maintaining network integrity and the tight regulation of network dynamics for pulsatile actomyosin and cell dynamics but do not allow us to distinguish between the roles of microtubule architecture and dynamics (Fig S3). Our experiments with EB1-DN however allowed us to specifically address the importance of persistent microtubule growth. Importantly, they allowed us to uncover a novel mode by which the microtubule cytoskeleton influences actomyosin dynamics that depends on the generation of moving patronin platforms that enable persistent microtubule growth. Our results differ from recent studies that have demonstrated an interplay between the two cytoskeletal systems (Booth et al., 2014; Ko et al., 2019) with respect to the nature of microtubule organisation (polarisation in the apical plane) or its consequences on constriction (regulation of pulse amplitude and constriction rate). Our results substantiate the role of microtubule plus ends as mobile platforms that can spatially regulate signalling and force generation through their interactions with other proteins (Akhmanova and Steinmetz, 2015; Rogers et al., 2004; Verma and Maresca, 2019). They identify the importance of RhoGEF2-EB1 interactions in the spatial regulation of Rho activity and actomyosin contractility. Recent work has also suggested the association between RhoGEF2 and EB1 in the ectoderm during Drosophila germ band extension (de las Bayonas et al., 2019). The comparable velocities and colocalization of RhoGEF2 and EB1 that we observe in real time suggests that RhoGEF2 tip-tracks. The transience of their association also suggests that RhoGEF2 is released from microtubule ends at the apical cortex. In this respect, our findings suggest a different mechanism than that employed during cytokinesis in Drosophila S2 cells in which the Pebble/Ect2 RhoGEF does not itself tip-track but depends on its interactions with the plus end association of centralspindlin to trigger local cortical contractility (Verma and Maresca, 2019). How the association or release of RhoGEF2 is influenced by microtubule dynamics or other + TIPs will be interesting to explore. Recent work has identified a role for patronin foci as non centrosomal microtubule organising centres that can nucleate microtubule arrays (Naschekin et al., 2016; Toya et al., 2016; Takeda et al., 2018). Our work uncovers mobile patronin platforms that can dynamically regulate the elaboration of microtubule cages and tails in subcellular space. What triggers their formation and drives their movement will be interesting to explore.

Another aspect that our findings uncovered is the regulation of the spatial distribution of actomyosin regulators along the radial axis by microtubule growth. That microtubule plus ends can modulate actomyosin contractility through their association with RhoGEF2 and other Rho regulators has been previously demonstrated (Verma and Maresca, 2019; Ito et al., 2017; Rafiq et al, 2019). The reversed radial polarity of Rho kinase (redistribution from the apicomedial pool to the circumapical pool) that we observed upon perturbing microtubule growth could rely on the increased availability of RhoGEF2 for cortical localisation. Alternatively, the two pools of active Rho could rely on different activators that are also differentially sensitive to microtubule dynamics (de las Bayonas et al., 2019).

Microtubules are capable of self-organising to form higher order network conformations in response to even weakly polarising external and internal cues by mechanical anisotropy (Bidhendi et al., 2019; Gomez et al., 2016; Meghana et al., 2011; Mirabet et al., 2018). Our analysis of the dynamics of apicomedial myosin and microtubule cages/asters suggests that the congregation of EB1 puncta and the condensation of patronin foci follow the appearance of a small myosin seed (Figure 4I). This suggests that actomyosin aggregates may break mechanical symmetry and trigger directional microtubule nucleation and growth. The work of Martin and colleagues (Ko et al., 2019) also revealed that the formation of medioapical patronin foci in ventral furrow cells depends on actomyosin contractility. Our results tempt the speculation that the subsequent dispersal of EB1 and patronin foci may also be influenced by the activation of contractility. Indeed, microtubule catastrophe has been demonstrated previously to occur upon contact with the cortex (Akhmanova and Steinmetz, 2008). Together, these findings suggest that microtubules may (re)act as first responders to anisotropies in tension. Whether and how microtubule growth and shrinkage are regulated by actomyosin contractility remains to be determined. (732)

## Acknowledgements

We thank Nick Brown, Damian Brunner, Jordan Raff, Tadashi Uemura, Kai Zinn, Melissa Rolls, the Bloomington Drosophila Stock Centre and the Developmental Studies Hybridoma Bank for reagents, Yusuke Toyama and Teng Xiang for the time integration code, Himanshu Bhat for the images in Figure S1B, members of the MN lab for discussion, Sudeepa Nandi for comments on the manuscript and TIFR/DAE (RT12001) for funds.

## Author contributions

AG, SS and DS designed and performed experiments, analysed data and contributed to manuscript writing. MN conceived and supervised the project, analysed data, wrote the manuscript and obtained funding.

## Materials and Methods

Key Resources Table.

| REAGENT or RESOURCE | SOURCE | IDENTIFIER |
| --- | --- | --- |
| **Antibodies** |  |  |
| DCAD2 (Anti ECadherin) | DSHB | AB_528120 |
| Anti GFP A11122 | Invitrogen | AB_221569 |
| Anti EB1 | Abcam | AB_53358 |
| pMLC Ser19 | Cell Signaling | Cell signaling_3671 |
| Anti RFP | Abcam | AB_945213 |
| Alpha tubulin | Sigma | AB_477593 |
| Acetylated tubulin | Santa Cruz | AB_628409 |
| Phalloidin 647 | Sigma | Sigma-Aldrich_65906 |
| DAPI | Sigma | Sigma-Aldrich_D-1388 |
| **Experimental Models: Organisms/Strains** |  |  |
| *D. melanogaster*:<br>Ubi::ECadherin GFP | Uemura et al., 1996.<br>Kind gift of Tadashi Uemura. | NA |
| *D. melanogaster*:<br>*y w sqh<sup>AX3</sup>*; Sqh GFP | Bloomington Drosophila<br>Stock Centre (BDSC) | BDSC_57144<br>Flybase_FBst0057144 |
| *D. melanogaster*:<br>EB1 GFP/ TM6B | Bulgakova et al., 2013.<br>Kind gift of Nick Brown. | NA |
| *D. melanogaster*:<br>Sqh mCherry/ CyO | Martin et al., 2009.<br>Kind gift of Aditya Saxena. | NA |
| *D. melanogaster*:<br>Ubi-p63E-Patronin<br>GFP/ CyO | Bloomington Drosophila<br>Stock Centre (BDSC) | BDSC_55128<br>Flybase_FBst0055128 |
| *D. melanogaster*:<br>Jupiter GFP | Buszczak et al., 2007. | NA |
| *D. melanogaster*:<br>UAS Tau | Bloomington Drosophila<br>Stock Centre (BDSC) | BDSC_51362<br>Flybase_FBst0051362 |
| *D. melanogaster*:<br>UAS Spastin GFP/TM3 | Jankovics and Brunner, 2006.<br>Kind gift of Damian Brunner | NA |
| *D. melanogaster*:<br>UAS EB1DN Cherry | Bulgakova et al., 2013.<br>Kind gift of Nick Brown | NA |
| *D. melanogaster*:<br>sqh::Sqh GFP | Bloomington Drosophila<br>Stock Centre (BDSC) | BDSC_57145<br>Flybase_FBst0057145 |
| *D. melanogaster*:<br>sqh::Rok GFP | Bloomington Drosophila<br>Stock Centre (BDSC) | BDSC_52289<br>Flybase_FBst0052289 |
| *D. melanogaster*:<br>sqh::RhoGEF2 GFP | Vienna Drosophila<br>Resource Centre (VDRC) | v318165<br>Flybase_FBst0491623 |
| *D. melanogaster*:<br>*w<sup>1118</sup>* | Bloomington Drosophila<br>Stock Centre (BDSC) | BDSC_5905<br>Flybase_FBst0005905 |
| *D. melanogaster*:<br>UAS-EB1 RFP | Mattie et al., 2010.<br>Kind gift of Melissa Rolls. | Flybase_FBtp0064377 |
| *D. melanogaster*:<br>Ubi-Spd2 GFP | Dix and Raff, 2007.<br>Kind gift of Jordan Raff. | NA |
| *D. melanogaster*:<br>c381 Gal4 | Bloomington Drosophila<br>Stock Centre (BDSC) | BDSC_3734<br>Flybase_FBst0003734 |
| **Software and Algorithms** |  |  |
| FIJI | NIH | SCR_002285 |
| Tissue Analyzer<br>(Fiji plugin) | Aigouy et al., 2016. | https://grr.gred-clermont.fr/labmirouse/software/WebPA |
| MTrack<br>(Fiji plugin) | Kapoor et al., 2019. | https://imagej.net/MicrotubuleTracker |
| Adobe Photoshop | Adobe | SCR_014199 |
| Adobe Illustrator | Adobe | SCR_010279 |
| RStudio |  | RRID:SCR_000432 |
| OriginPro |  | RRID:SCR_014212 |
| BoxPlotR |  | RRID:SCR_015629 |
| GraphPad | GraphPad | http://www.graphpad.com |

### Drosophila stocks

The following transgenic lines were used: c381Gal4 (AS Gal4, for driving expression in the amnioserosa), Jupiter GFP [to mark microtubules; (Buszczak et al., 2007)], Ubi::Patronin GFP (to mark the minus ends of microtubules, Wang et al, 2013), Sqh mCherry and Sqh GFP (to mark non muscle myosin II), sqh::Rok GFP (Rho kinase tagged to GFP; Abreu-Blanco et al., 2014), RhoGEF2 GFP (MiMIC line, Sarov et al., 2016) and sqh::RhoGEF2 GFP (RhoGEF2 tagged to GFP; Nakamura et al., 2017) were obtained from the Bloomington Drosophila stock Centre. EB1 GFP (to mark microtubule plus ends), and UAS EB1-DN Cherry (to perturb the persistent growth of microtubules) were kind gifts from Nick Brown, Cambridge, UK (Bulgakova et al., 2013), UAS-EB1 RFP (Mattie et al., 2010) was a kind gift of Mellisa Rolls, Ubi::ECadh GFP (to mark adherens junctions was a kind gift of Tadashi Uemura (Uemura et al., 1996). UAS Tau (from the Bloomington Drosophila Stock Centre) was used to cause microtubule bundling and increase stability (Chatterjee et al., 2009), and UAS Spastin /UAS Spastin GFP was used to sever microtubules [(Jankovics and Brunner, 2006; Sherwood et al., 2004); kind gifts of Kai Zinn, Caltech and Damian Brunner, EMBL]. Ubi::Spd2 GFP was a kind gift from Jordan Raff (Dix and Raff, 2007).

### Genotypes analysed

#### Figure 1

A1-A7”-*w*; Sqh mCherry/ Ubi::Patronin GFP was used to visualize myosin along with microtubule minus end dynamics.

B1-B7” & C1-C6’-*w*; Sqh mCherry/+; EB1 GFP/ + was used to visualize myosin along with microtubule plus end dynamics.

D1-D6 – w; Ubi::Patronin GFP was used for immunofluorescence experiments to examine the relationship between microtubule ends and myosin.

#### Figure 2

*w*; Ubi::ECadherin GFP/+, c381 Gal4/+

*w*; Ubi::ECadherin GFP/+; UAS EB1-DN Cherry/+; c381 Gal4/+

Both genotypes were used to quantify pulse amplitude, cell area dynamics, tissue dynamics and frequency of delamination.

#### Figure 3

A1-A7 & C-G-*w*; Sqh GFP;; c381 Gal4/+

B1-B7 & C-G -*w*; Sqh GFP/+; UAS EB1-DN Cherry/+; c381 Gal4/+

Both genotypes were used to visualize and quantify myosin dynamics in control embryos and in embryos expressing the dominant negative of EB1 in the amnioserosa.

#### Figure 4

A1-A5-*w*; sqh::RhoGEF2 GFP was used to visualize RhoGEF2 dynamics.

B1-B4-*w*;; RhoGEF2 GFP was used for immunofluorescence experiments to examine the relationship between microtubules, actin and RhoGEF2.

C1-C5-sqh::RhoGEF2 GFP/+; UAS EB1 RFP/+; c381 Gal4/+ was used to examine the dynamics of RhoGEF2 with respect to microtubule plus ends.

D1-D7 and F1-F1’-*w*; +/+; sqh::Rok GFP/+; c381 Gal4/+ was used in immunofluorescence experiments and to examine the dynamics of Rho kinase.

E1-E7 and F2-F2’-*w*; +/+; sqh::Rok GFP/UAS EB1-DN Cherry; c381 Gal4/+ was used in immunofluorescence experiments and to examine the dynamics of Rho kinase upon overexpressing dominant negative form of EB1 in the amnioserosa.

#### Figure S1

A1-A4’-*w*^*1118*^ was used in immunofluorescence experiments to examine the organisation of microtubule and actin.

B1-B3-Ubi-Spd2 GFP was used to do examine the organisation of the microtubule network with respect to centrosome using immunofluorescence.

C1-C3 and C1’-C3’-*w*^*1118*^ was used in immunofluorescence experiments to examine the distribution of acetylated tubulin.

D-D’-*w*^*1118*^

E-E’-*w*; +/+; UAS Spastin GFP/+; c381 Gal4/+

F-F’-*w*; UAS Tau/+; +/+; c381 Gal4/+

G-G’-*w*; +/+; UAS EB1-DN Cherry/+; c381 Gal4/+

All the above genotypes (D-G) were used to look at the microtubule network using immunofluorescence.

#### Figure S2

A1-A7: *w*;; Sqh GFP was used to look at myosin dynamics.

B1-B6-*w*; Ubi::Patronin GFP was used to look at microtubule minus end dynamics.

C1-C6-*w*;; EB1 GFP/ TM6 was used to look at microtubule plus end dynamics.

D1-D6-*w*; Ubi::ECadherin GFP/+; EB1 GFP/ + was used to look apical cell membrane and microtubule plus end dynamics.

E1-E6 and F1-F5-*w*;; Jupiter GFP was used to visualize microtubule dynamics.

G1-G5-*w*; Ubi::ECadherin GFP/+; Jupiter GFP/+, was used to look apical cell membrane and microtubule dynamics.

H1-H7”-*w*; Sqh mCherry/+; Jupiter GFP/+ was used to visualize myosin and microtubule dynamics.

#### Figure S3

A1-A7 & D-H-*w*; Sqh GFP;; c381 Gal4/+ was used to visualize and quantify myosin dynamics in wild type embryos.

B1-B7 & D-H -*w*; Sqh GFP/+; UAS Spastin GFP/+; c381Gal4/+ was used to visualize and quantify myosin dynamics upon overexpressing spastin in the amnioserosa.

C1-C7 & D-H - *w*; Sqh GFP UAS Tau/+; +/+; c381Gal4/+ was used to visualize and quantify myosin dynamics upon overexpressing Tau in the amnioserosa.

### Embryo harvesting and Imaging

For live imaging, flies were allowed to lay for 4 hours, and embryos aged at 29°C for 8 hours to enrich for stages of dorsal closure. They were then dechorionated using 50% bleach for 2 minutes and washed with water. Embryos were then put on a 0.5 mm coverslip (Corning) on a thin film of Halocarbon oil 700 (Sigma) and imaged on an inverted microscope (Olympus FluoView 1000 confocal microscope). For fixed preparations, embryos were harvested and stained using standard protocols (Narasimha and Brown, 2006). The following primary antibodies were used: anti-GFP (rabbit 1:1000, Invitrogen), anti-EB1 (rat 1:200, Abcam), pMLC Ser19 (rabbit 1:50, Cell signaling) anti-RFP (mouse 1:1000, Abcam), anti-ECadherin (rat 1:10; DSHB), anti-alpha tubulin (mouse 1:1000, Abcam), and anti-acetylated tubulin (mouse 1:500, Sigma). Alexa Fluor conjugated secondary antibodies (Invitrogen) were used at 1:200 dilutions, and Phalloidin Atto 647 (Sigma) was used at 1:100 dilution. Dapi was used to label nuclei. Embryos were then stored and mounted in Vectashield and confocal images were taken on Olympus FluoView 1000 confocal microscope.

### Quantitative morphodynamics

Apical areas of individual cells in Phase I of dorsal closure were measured for a duration of 140 minutes prior to the closure, from maximum intensity projected, time-lapse images of E-Cadherin GFP. Central amnioserosa cells were segmented using Tissue analyzer (Aigouy et al., 2016), the selected ROIs exported to Fiji (Schindelin et al., 2012), and the areas extracted. Cell area dynamics was analysed by plotting the changes in apical area as a function of time (Microsoft Excel). Mean rates of change of area (± sd) and the linear regression was done using Origin Pro 2015. Similarly obtained cell area traces were also used to measure the amplitudes of pulses (Saravanan et al, 2013). Normalized pulse amplitude was calculated as (*A*_max_/*A*_0_) − (*A*_min_/*A*_0_), where *A*_max_ and *A*_min_ represent, respectively, the maximum and minimum apical cell areas during a pulse and *A*_0_ denotes the initial cell area. Pulse amplitude was calculated from pulses observed in cells in Phase I over a 30-minute interval and averaged over multiple cells from multiple embryos of the same genotype to obtain mean ± range. The area of the amnioserosa was measured as the area of the ellipse bound by the leading edge/actin cable from 120 minutes prior to the completion of closure to the end of closure, from maximum intensity projected, time-lapse images of E-Cadherin GFP. The areas were extracted and plotted as mentioned above. The number of cell delamination events occurring in the time window from the formation of the posterior canthus formation to the end of dorsal closure were marked and counted using Fiji. The significance of differences between means of different genotypes was calculated using unpaired Student’s t test on GraphPad (Prism Software).

### Quantification of myosin dynamics

Formation time of the myosin blob was defined as the time taken for the attainment of its largest size from the point of its initial detection. Dissolution time was defined as the time taken by the largest blob to disappear completely. Total persistence time was obtained by adding the formation and dissolution times. Myosin times from several myosin cycles from multiple central amnioserosa cells in Phase I of dorsal closure from different embryos of the same genotype were pooled to obtain the mean myosin cycle times.

Path length was defined as the distance travelled by the coalesced myosin blob, whose trajectory was tracked manually using an ImageJ plugin MTrack (Kapoor et al., 2019), from the time it was first detected till it disappeared. The plugin also marks the traced tracks to enable the visualization of qualitative differences. The path lengths of several myosin cycles from multiple central amnioserosa cells in Phase I from different embryos of the same genotype were pooled to obtain the mean path length for each genotype. The path length of each myosin blob was divided by its respective persistence time to obtain its velocity and the mean velocity was determined from the trajectories of multiple myosin blobs. The significance of differences between the means obtained for different genotypes was determined using the unpaired Student’s t test on GraphPad (Prism Software).

### Velocity measurements for EB1 and RhoGEF2

Movies of EB1 GFP and sqh::RhoGEF2 GFP were used to calculate the velocity of EB1 comets and RhoGEF2 comets respectively. The distance travelled by the comets was calculated manually using Fiji by tracking the tip of the comet in one direction till it disappeared or began to retract. This was divided by the time taken to travel the measured distance to obtain the velocity. The significance of differences between the mean velocities of the two types of comets was determined using the unpaired Student’s t test on Origin Pro.

### Time integrated EB1 tracks

As the EB1 comets are highly dynamic, we used a time integration macro (written and kindly provided by Teng Xiang and Yusuke Toyama, Mechanobiology Institute, Singapore) that integrates the dynamics of three successive frames, to obtain a better visualisation of their trajectories.

### Polar plots

Polar plots were used to get a better representation of the spatio-temporal relationship between myosin and microtubule ends (Patronin and EB1). The images were subjected to background subtraction using median filter, followed by thresholding and identification of objects using Analyze particles in Fiji. The X and Y coordinates of the extracted particles were used to calculate the distance ‘r’ and the angle ‘theta’ from the center of the circle (ROI marked in the corresponding images) using RStudio. Polar plots were then plotted using OriginPro. Size of the circle was made proportional to the area occupied by the objects.

### Intensity measurements

Rok intensity was measured from maximum intensity projections of apical slices of the central amnioserosa cells during Phase I of dorsal closure from embryos carrying sqh::Rok-GFP that had been stained with an anti-GFP antibody. Straight lines were drawn from one interface of the cell to the opposite interface through the medial blob or the diffuse medial pool of Rok. Line intensity profiles were obtained using Fiji (Schindelin et al., 2012). The intensity profile along the normalised length (from 0-1, with 0 and 1 being at the two membranes) was then plotted in Microsoft Excel.

### Graphs and statistical analysis

All statistical analysis was done using GraphPad Prism software. Box plots were made using BoxPlotR (Spitzer et al., 2014) and show median ± interquartile range. Thick black lines show the median, the plus mark shows the mean, and the box limits indicate 25^th^ and 75^th^ percentile. Whiskers extend to 1.5 times the interquartile range from the 25^th^ and 75^th^ percentiles. Individual data points are represented by dots. The sample size (n) analysed for the individual genotypes is mentioned in the graph. Unpaired Student’s t-test was used to assess the significance of difference between the means for all the statistical analysis.

## Supplementary Figure Legends

**Figure S1:**
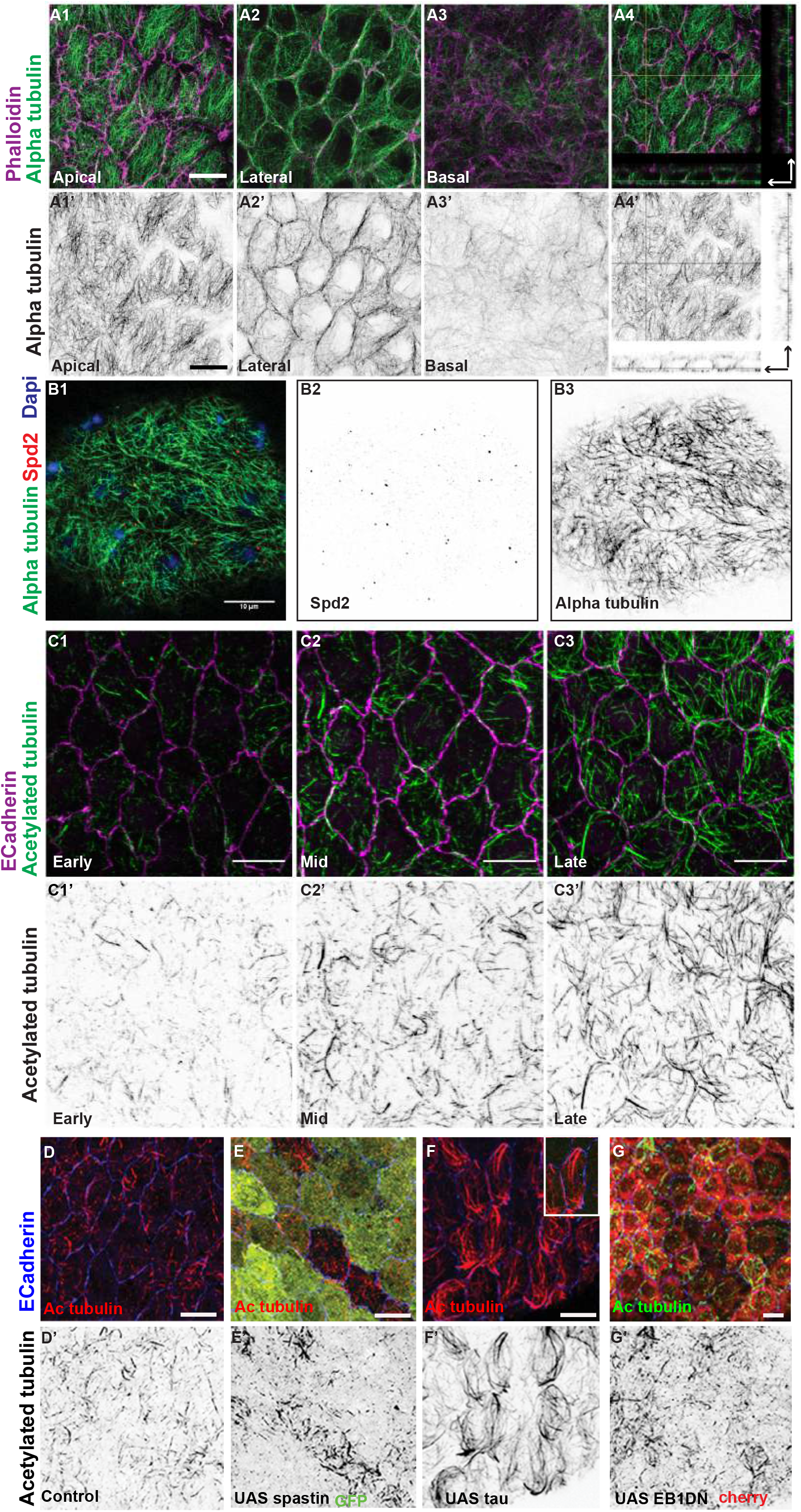
Organisation of the microtubule cytoskeleton in the amnioserosa. Apical (A1), lateral (A2), basal (A3) and orthogonal (A4) sections of amnioserosa cells during mid dorsal closure, stained with alpha tubulin (green) and phalloidin (magenta). A1’-A4’ show the inverted grey scale images of tubulin from the merged images in A1-A4. (B) The microtubule network visualised using alpha tubulin (green in B1, grey in B3) in an embryo expressing the centrosomal protein Spd2 tagged to GFP (red in B1 and grey in B2). The apical meshwork does not show a radial pattern emanating from the centrosomes labelled with Spd2. The nuclei are labelled with DAPI (blue). (C1-C3) The stable pool of apical microtubules visualised using acetylated tubulin (green) during early (C1), mid (C2) and late (C3) stages of dorsal closure. (D-G) Apical microtubule network visualised using acetylated tubulin (red in D-F and green in G) in wild type embryos (D) and embryos overexpressing Spastin (E), Tau (F) and EB1-DN (G). C1’-C3’ and D’-G’ shows the corresponding single channel images for acetylated tubulin. Cell boundaries are marked with ECadherin (magenta in C1-C3 and blue in D-G). Scale bar-10μm. (Related to Figure 1)

**Figure S2:**
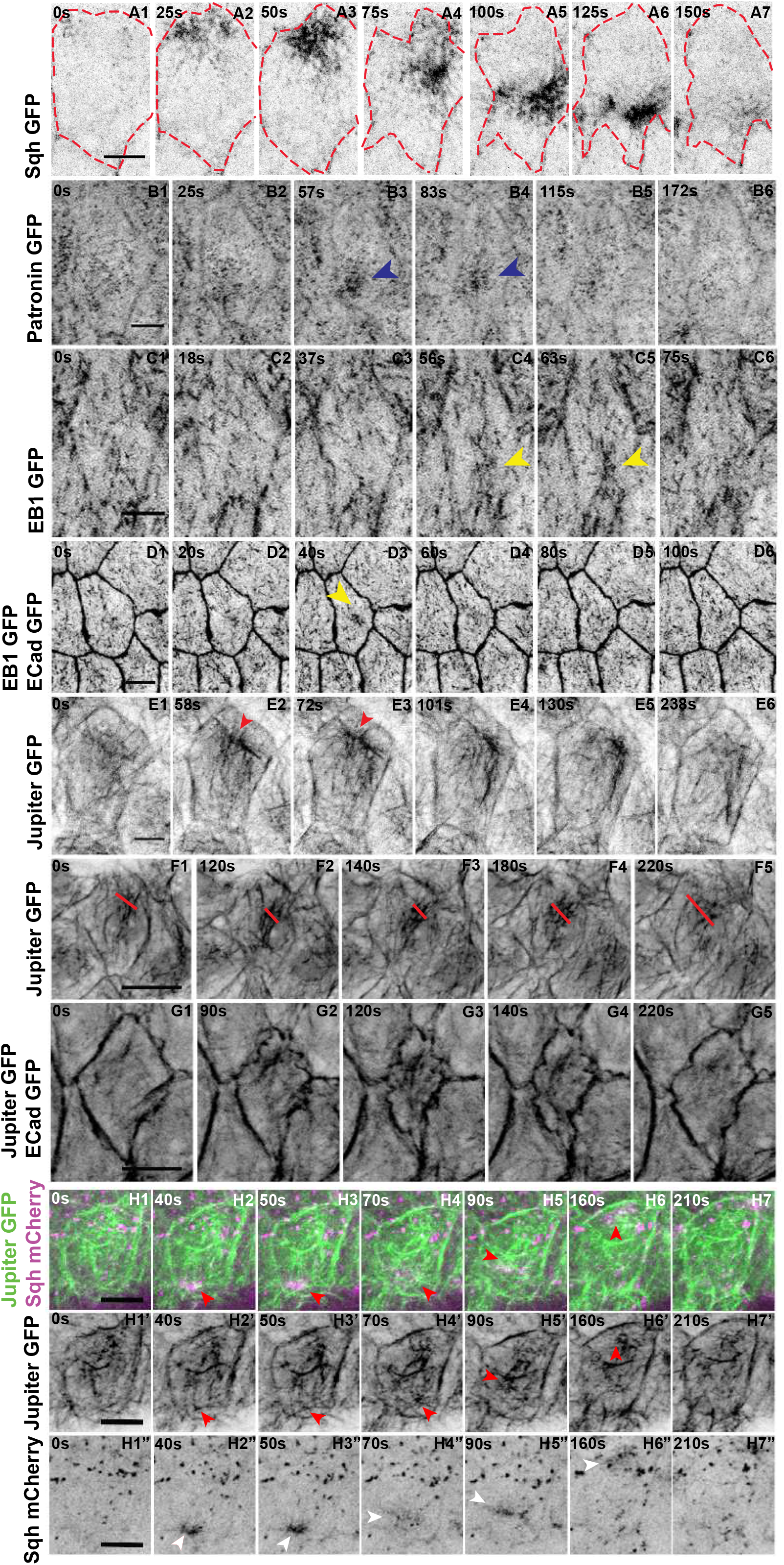
Dynamics of myosin and microtubules during pulsed apical constriction. (A1-A7) Time-lapse images of an amnioserosa cell in Phase I of dorsal closure showing the dynamic changes in the organisation of myosin (visualised with Sqh GFP, grey/black) during a single pulse. The dashed red line marks the apical cell membrane. (B1-B6) Time-lapse images of an amnioserosa cell in Phase I of dorsal closure, showing the dynamics of microtubule minus ends (Patronin GFP) during a single pulse. Blue arrowheads in B3, B4 point to the condensing patronin cloud. (C1-C6) Time-lapse images of amnioserosa cells in Phase I of dorsal closure, showing the congregation (yellow arrowheads) and dispersal of microtubule plus ends (EB1 GFP comets) during a single pulse. (D1-D6) Temporal correlation between the congregation of EB1 GFP comets and apical cell area dynamics (ECadherin GFP). (E1-E6, F1-F5) Time-lapse images of amnioserosa cells in Phase I of dorsal closure, showing the dynamic changes in the organisation of the apical microtubule meshwork (Jupiter GFP, grey/black), showing the formation and dispersal of microtubule aster-like configurations during a pulse (red arrowheads in E2, E3), and the ‘bundling” and splaying of microtubules (red lines in F1-F5). (G1-G5) Temporal correlation between microtubule “bundling” (Jupiter GFP) and apical cell area dynamics (ECadherin GFP) during a single pulse. (H1-H7) Time-lapse images of an amnioserosa cell in Phase I of dorsal closure from embryos carrying Jupiter GFP (green) and Sqh mCherry (magenta), showing the reorganization of the apical microtubule meshwork (white arrowheads) around a medial myosin blob during a single pulse. H1’-H7’ and H1”-H7” are the corresponding single channel images of Jupiter GFP and Sqh mCherry respectively. Red and white arrowheads point to the microtubule asters and the myosin blob respectively. Scale bar-5 μm. (Related to Figure 1. See also Videos S2, S3.)

**Figure S3:**
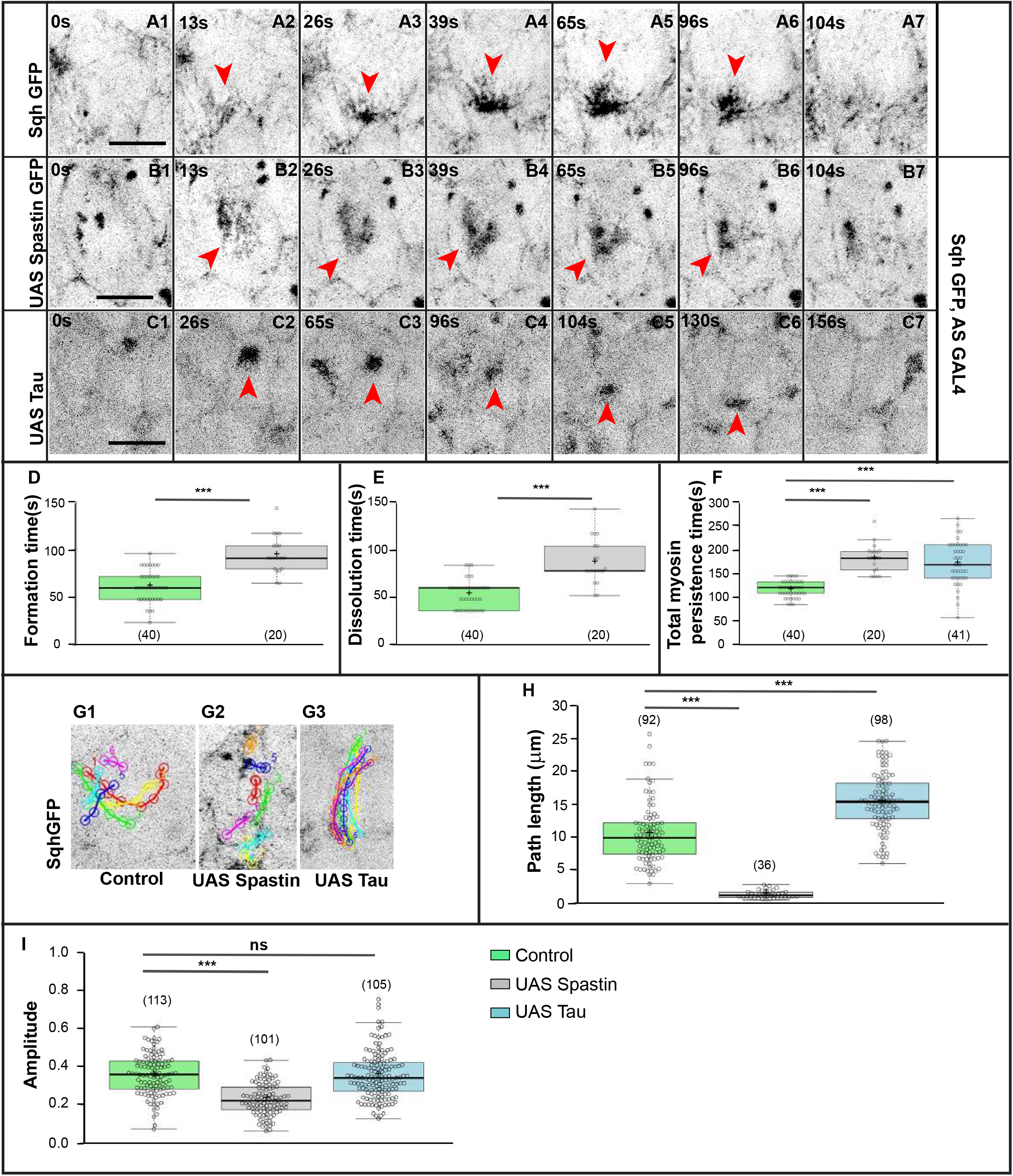
The integrity and dynamics of the microtubule cytoskeleton influences apicomedial myosin organisation and dynamics. (A-C) Time-lapse images of single amnioserosa cells during Phase I of dorsal closure showing qualitative changes in the spatial organisation of apicomedial myosin (visualised using Sqh GFP) in embryos that are otherwise wild type ([A1-A7; (n=40 myosin cycles from 4 embryos)], or overexpress Spastin [B1-B7; (n=20 myosin cycles from 3 embryos)], or tau [C1-C7; (n=41myosin cycles from 4 embryos)] in the amnioserosa. Red arrowheads point to apicomedial myosin complexes. Scale bar −10μm. (D-F) Myosin cycle times (Formation-D, dissolution-E and total myosin persistence times-F) measured in wildtype embryos, and in embryos overexpressing Spastin or Tau in a Sqh GFP /+ background. (G) Representative tracks of medial myosin movement in a single Phase I amnioserosa cell from control embryos (G1), and from embryos overexpressing UAS Spastin (G2) or UAS Tau (G3). Each track represents the distance travelled by one apicomedial myosin blob/structure in one cycle. Many myosin cycles (colour coded) were tracked in each cell. (H) Quantitative analysis of apicomedial myosin path lengths in each of the genotypes analysed. (I) Normalised pulse amplitude of amnioserosa cells from control embryos and embryos overexpressing Spastin or Tau in the amnioserosa during phase I (pulsed constriction) of dorsal closure (see Methods). In the Box plots in D-F, H, I boxes show median (dark line) ± interquartile range. The mean is also indicated by ‘+’. The sample size is given in brackets. ***-p<0.0001. (Related to Figure 3.)

## Movie Legends

**Video S1: Microtubule ends form cages and tails around apicomedial myosin.**

A) Microtubule minus ends are visualized by Patronin GFP and myosin by Sqh mCherry. B, C) Microtubule plus ends visualized by EB1 GFP (time integrated tracks in C) show a transient association with apicomedial myosin visualized by Sqh mCherry. Red arrowheads indicate the myosin blob and yellow arrowheads indicate the reorganising microtubule ends. Scale bar – 5μm. (Related to Figure 1.)

**Video S2: Apical microtubule network reorganisation during pulsed apical constriction; cages/asters and bundles.**

A, B) The apical microtubule cytoskeleton visualized using Jupiter GFP showing its reorganisation into cages/ aster-like configurations (A, red arrowheads), microtubule bundles (B, yellow line) or buckling (B, yellow arrowhead). C) The association of microtubules (green) with myosin (magenta). Red arrowheads indicate asters/cages and white arrowheads indicate myosin. Bundled microtubules can also be seen. Scale bar - 5 μm. (Related to Figure S2.)

**Video S3: Microtubule ends reorganize during pulsed apical constriction.**

(A, B) Microtubule plus ends visualized by EB1 GFP congregate and disperse during the constriction and relaxation phases of a pulse respectively. Microtubule minus ends visualized by Patronin GFP form a dense cloud that condenses as the cells constrict (C). Red arrowheads mark the reorganisation. Cell outlines are marked with E-Cadherin GFP in B and by red lines in A, C. Scale bar - 5 μm unless specified. (Related to Figure S2.)

**Video S4: Microtubule growth modulates cytoskeletal, cell and tissue dynamics in the amnioserosa.**

A) Myosin dynamics visualized using Sqh GFP in control and EB1-DN expressing amnioserosa cells. Coloured traces represent tracks of individual myosin blobs (black). B) Cell delamination (blue dots) and contraction of the amnioserosa visualized using ECadh GFP in control and EB1-DN expressing embryos. Scale bar - 5 μm unless specified. (Related to Figures 2 and 3.)

**Video S5: RhoGEF2-EB1 interactions govern the spatial distribution of Rho pathway activity.**

A) The dynamics of RhoGEF2 in Phase 1 amnioserosa cells. The red outline marks the cell boundary and red arrowheads point to the congregation, movement and dispersal of RhoGEF2 clusters. B) The transient association (within the yellow boxes) of RhoGEF2 (green) with EB1 (magenta) in Phase 1 amnioserosa cells. C) The localisation and dynamics of Rho kinase (Rok) control and in EB1-DN amnisoerosa cells (right) overexpressing cells. The yellow arrowhead indicates the medial Rok blob and the red arrowhead indicates circumapical Rok. Scale bar - 5 μm. (Related to Figure 4.)

